# Neural correlates of beauty retouching to enhance attractiveness of self-depictions in women

**DOI:** 10.1101/2020.12.25.424237

**Authors:** Chisa Ota, Tamami Nakano

## Abstract

Beauty filters, while often employed for retouching photos to appear more attractive on social media, when used in excess cause images to give a distorted impression. The neural mechanisms underlying this change in facial attractiveness according to beauty retouching level remain unknown. The present study used functional magnetic resonance imaging in women as they viewed photos of their own face or unknown faces that had been retouched at three levels: no, mild, and extreme. The activity in the nucleus accumbens (NA) exhibited a positive correlation with facial attractiveness, whereas amygdala activity showed a negative correlation with attractiveness. Even though the participants rated others’ faces as more attractive than their own, the NA showed increased activity only for their mildly retouched own face and the amygdala exhibited greater activation in the others’ faces condition than the own face condition. Moreover, amygdala activity was greater for extremely retouched faces than for unretouched or mildly retouched faces for both conditions. Frontotemporal and cortical midline areas showed greater activation for one’s own than others’ faces, but such self-related activation was absent when extremely retouched. These results suggest that neural activity dynamically switches between the NA and amygdala according to perceived attractiveness of one’s face.

## Introduction

In Renaissance Italy, ladies made their eyes seem more alluring by inducing pupillary mydriasis with applied drops of belladonna extract (Forbes, 1977). About 500 years later, modern young women are retouching their self-portraits to make their eyes bigger and are actively posting them on the social network services. The woman’s desire to look more beautiful seems to be unbending regardless of time. Moderate beauty retouching can increase facial attractiveness, but in cases of excessive use it can make a face appear unnatural and even creepy. Ironically, the facial attractiveness then becomes much lower than that of the original face (Nakano & Uesugi, 2020). Since extreme facial retouching creates a face whose balance between individual parts and overall structure deviates from the distribution of real humans (e.g. extremely big eyes and very thin chins), the negative impression towards the extremely retouched faces may be related to the “uncanny valley” phenomenon, in which eerie sensations are induced when viewing subjects with slight differences that cause them to look like false or unreal humans (Feng et al., 2018; Mori, MacDorman, & Kageki, 2012). Like this, facial impression dramatically and non-linearly changes depending on the degree of retouching, yet it remains unknown what neural mechanisms are involved in this inverted U-shaped change in perceived facial attractiveness.

Previous brain imaging studies consistently reported that attractive faces induced activation in reward circuits. Aharon et al. (2001) showed that beautiful female faces activate the nucleus accumbens (NA) of male brains. This NA, receiving the dopaminergic input from the ventral tegmental area (VTA), is the center of motivation, reward, and reinforcement learning (Covey & Cheer, 2019). Another study also revealed that attractive faces induced brain activation in the orbital-frontal cortex (OFC) and medial prefrontal cortex (mPFC), regions involved in representing stimulus reward value (O’Doherty et al., 2003). Additionally, enhancing facial attractiveness by using makeup increased neural activity in the OFC and medial prefrontal regions (Ueno et al., 2014). Considering this evidence, it can be presumed that the reward system of the brain plays a central role in the positive effect of beauty retouching perceived face attractiveness.

On the other hand, the neural substrates underlying the negative effects of beauty retouching remains unclear. One candidate brain region is the amygdala, which plays a central role in processing negative facial information and aversive learning (Adolphs, Russell, & Tranel, 1999; LeDoux, 2000; Schienle, Schafer, Stark, Walter, & Vaitl, 2005; Vuilleumier, Richardson, Armony, Driver, & Dolan, 2004). A previous study reported that faces with low attractiveness induced greater activation in the amygdala than faces with moderate attractiveness (Winston, O’Doherty, Kilner, Perrett, & Dolan, 2007). Moreover, the “Thatcher illusion“, in which a transfigured face with inverted eyes and mouth induces a strong eerie impression (Thompson, 1980), elicits strong activation in the bilateral amygdala (Rotshtein, Malach, Hadar, Graif, & Hendler, 2001). This evidence indicates that the amygdala is involved in the decrease in perceived facial attractiveness and eerie sensation felt towards faces with an extreme beauty retouching applied. Considering these positive and negative effects of beauty retouching, we speculated that the inverted U-shaped change in perceived facial attractiveness based on beauty retouching level is caused by the interplay between positive valence systems, which code beautiful faces as rewards, and negative valence systems, which code deviated human faces as aversive stimuli.

We should also consider that this interaction between two opposite valence systems may work differently between cases of own face versus unknown face because self-related information is specially processed in our brain. A meta-analysis of neuroimaging studies revealed that self-related processing activates medial cortical regions (Northoff et al., 2006) that overlap with the brain regions showing activation in response to facial attractiveness (O’Doherty et al., 2003; Winston et al., 2007). A previous study, which manipulated facial attractiveness by contrast effect, also reported that medial cortical regions and the VTA, the center of human dopaminergic reward system, exhibit greater activation when self-face looks more attractive than other-face (Oikawa et al., 2012). Consistently, our previous behavioral study has shown that young women prefer stronger beauty filters on their own face than on others’ faces (Nakano & Uesugi, 2020). This suggests that the enhanced attractiveness of self-face has a higher reward value than that for other-face and that, as a result, the preference for using beauty filters on their own face is reinforced. This evidence raises the possibility that positive valence systems work only for enhanced attractiveness of self-face and not for that of other-face. Rather, the amygdala may exhibit neural activity in response to face with extreme filter and unknown face. To address this possibility, we compared the neural activity, which reflects the change in facial attractiveness according to the level of beauty filter applied (no filter, mild filter, extreme filter), between self-face and other-face conditions through functional magnetic resonance imaging (fMRI). This study focused on women because retouching behavior is popular among women to depict themselves more beautiful (Kee & Farid, 2011).

## Methods

### Participants

Thirty-three females participated in this study (mean age: 21.6 years ranging from 19 to 24 years). They had no abnormal neurological history and had normal vision either uncorrected or corrected by glasses. The review board of Osaka University approved the experimental protocol (FBS30-4), and our procedures followed the guidelines outlined by the Declaration of Helsinki. All participants provided written, informed consent prior to the experiment.

### Stimuli and Experimental Procedure

Before the fMRI experiment, we took a picture of each participant’s face as they stood in front of a white background with a black cape across their shoulders. For each participant, we took pictures of 10 different facial expressions without emotion (frontal face, face uttering “a”, “i”, “u”, “e”, “o”, and faces tilted to the right, left, up and down). These photos were used as the self-face stimuli. In addition, we took pictures of 10 women (mean age: 22.3 years, 20 to 25 years old), who were not acquainted with the fMRI participants, for use as other-face stimuli. Next, a beauty retouching application (free software SNOW, Snow Corp.) that enlarges the eyes and makes the chin smaller was applied to these photos in two stages: mild and extreme (Fig. 1A). For each participant, we prepared a series of self-face and other-face images with either no retouched, a mildly retouched, or an extremely retouched. All photo images were 600×750 pixels in size and converted to gray scale.

During the experiment, the participants laid in an MRI scanner while wearing earplugs and immobilizing their heads using sponge cushions, and viewed visual stimuli on a screen (1920×1080 pixel, viewing angle = 27.1 degrees) via a mirror placed in front of their eyes. In each trial, following the presentation of the fixation cross for 4-6 s, the photo of a face was presented for 3 s with gray background (Fig. 1B). Then, the participants were asked to rate attractiveness of the face on a scale of 1 (unattractive) to 8 (very attractive) within 3 s using two MRI-compatible four-button response devices placed under their hands. Ten trials were presented for each face type, for a total of 60 trials in one session. The order of the face stimuli was randomized across the participants. All participants completed two sessions, with a short break between sessions, for a total of 120 trials. Different facial images were used in each session.

We also asked three additional females who did not enroll in the fMRI experiment to evaluate facial attractiveness of all face stimuli with a scale of 1 to 8. The mean attractiveness score of self-face stimuli was 4.0 and that of other-face stimuli was 4.4. In each evaluator, there was no significant difference in facial attractiveness between self-face and other-face stimuli (two sample t-test).

### Data Acquisition

Functional images were acquired using multi-band T2*-weighted gradient echo type echo-planar imaging (EPI) sequences, which were obtained using a 3-Tesla MRI scanner (MAGNETOM Vida, Siemens) with a 64-channel array coil. We collected 750 scans per session (slice number = 45, slice thickness = 3 mm, repetition time [TR] = 1,000 ms, echo time [TE] = 30 ms, flip angle = 60 degrees, field of view [FOV] = 192×192 mm, voxel size [x, y, z] = 3×3×3 mm, multiband factor = 3). For an anatomical reference image, a T1-weighted structural image was acquired for each subject (MP-RAGE sequence, slice thickness = 1 mm, repetition time [TR] = 1,900 ms, echo time [TE] = 3.37 ms, flip angle = 9 degrees, field of view [FOV] = 256×256 mm, voxel size [x, y, z] = 1×1×1 mm). To correct the geometric distortion in EPI, we also acquired field maps for each participant (Siemens standard double echo gradient echo fieldmap sequence, slice thickness = 3 mm, repetition time [TR] = 753 ms, echo time [TE] = 5.16 ms, flip angle = 90 degrees, field of view [FOV] = 192×192 mm, voxel size [x, y, z] = 3×3×3 mm).

### Imaging Data Analysis

Acquired MRI data was processed using SPM12 and MATLAB 2019a. We discarded the first 3 EPI images in each session. To correct image distortion due to field inhomogeneity, field map correction was applied using the SPM fieldmap toolbox. We confirmed that the head movements were less than 3 mm in all participants, and the EPI images were realigned and unwarped. Each participant’s structural image was coregistered to the mean of the motion-corrected functional images. Then the EPI images were normalized to the standard brain template (Montréal Neurological Institute template), and smoothed using a Gaussian kernel filter with an 8-mm full-width-at-half-maximum (FWHM). We further checked signal drop-out of the EPI images of all participants by superimposing the section images of the implicit mask on the structural image (see the supplementary figure 1). Despite the field map correction, a part of the ventral frontal and anterior temporal lobes was still uncovered. Thus, the activity of these uncovered brain regions is unknown in the present study.

After pre-processing, we conduct a voxel-by-voxel regression analysis of expected hemodynamic changes for the 6 conditions (self/other × filter level) using the general linear model (GLM). The design matrix consisted of 6 conditions of face stimuli and button press, 6 realignment parameters per session. Data from one session was excluded for 3 participants due to sleeping (n=2) and an experimental error (n=1). Next, we conducted a parametric modulation analysis using attractiveness score as the parameter. For whole brain analyses, we used a family wise error rate (FWE) cluster-corrected threshold of p < 0.05 using a cluster-defining threshold of p < 0.001. Based on prior work (Motoki et al., 2016; Rotshtein et al., 2001; Schwartz et al., 2003; Whalen et al., 2004), we had a priori hypotheses that amygdala is involved in negative impression toward face images. Therefore, we used a small-volume corrected FWE cluster-level threshold of p < 0.05 in spheres of 8 mm around previous coordinates in the amygdala ([±27, −4, −20]) (Motoki et al., 2016).

To compare the temporal dynamics of the BOLD signal changes in the Region of Interest (ROI) between conditions, we extracted time course data of signal intensity in each ROI for each participant, applied a 128 Hz high-pass filter and linearly interpolated at 0.1 s resolution, and converted to z-score using mean and variance. Then, the time course was averaged for each condition across trials from 2 s before to 15 s after the face’s onset. The averaged beta value for each ROI was also calculated and compared between conditions using a two-way Analysis of variance (ANOVA) with factors of face type (self vs other) and retouching level (no, mild and extreme).

We also analyzed directed functional connectivity between ROIs by the multivariate Granger causality analysis concerning time course of signal intensity in each ROI using the MVGC toolbox (Barnett and Seth, 2014) with MATLAB. For analyzing the causal relationship between ROIs, we selected 5 ROIs that showed varied neural activation depending on facial attractiveness. In this analysis, time series data was resampled every 1 sec and model order was set to one. The Granger causality (GC) values were calculated across all epochs (each epoch was 15 seconds after the face onset) in each participant. The baseline GC values were also calculated (5 seconds before the face onset). Then, we detected the causal connection which significantly increased in the GC value against baseline using the paired t-test (alpha = 0.05; multiple comparisons were corrected using Bonferroni method).

## Results

### Behavioral results

First, we analyzed the effects of the beauty filter on facial impressions. As shown in Figure 2, the participants evaluated the faces with mild filter as the most attractive (self 4.16 ± 1.0; other 4.97 ± 0.79, mean ± s.d.) and the faces with extreme filter as the least attractive for both self and other-faces (self 2.09 ± 1.06; other 3.16 ± 1.07). The original faces without any filter were rated slightly less attractive than the faces with the mild filter but much more attractive than the faces with extreme filter (self 3.53 ± 1.01; other 4.36 ± 0.79). Across all filter levels, the mean attractiveness score of the other-face was much higher than that of the self-face. Two-way Analysis of Variance (ANOVA) with factors of face type (self/other) and retouching level (none, mild, extreme) revealed a significant main effect from the face type (F_1,32_ = 56.4, p < 0.00001) and the retouching level (F_2,64_ = 55.6, p < 0.00001), but no significant interaction between them (F_2,64_ =1.8, p = 0.17). The post-hoc analysis using the Ryan method confirmed that the faces with a mild retouching were significantly more attractive than both the faces with an extreme retouching (t_64_ = 10.3, p < 0.00001) and the original faces with no retouching (t_64_ = 3.3, p < 0.002). In addition, the faces with an extreme retouching were significantly less attractive than their corresponding original face (t_64_ = 7.0, p < 0.00001).

**Fig. 2.**
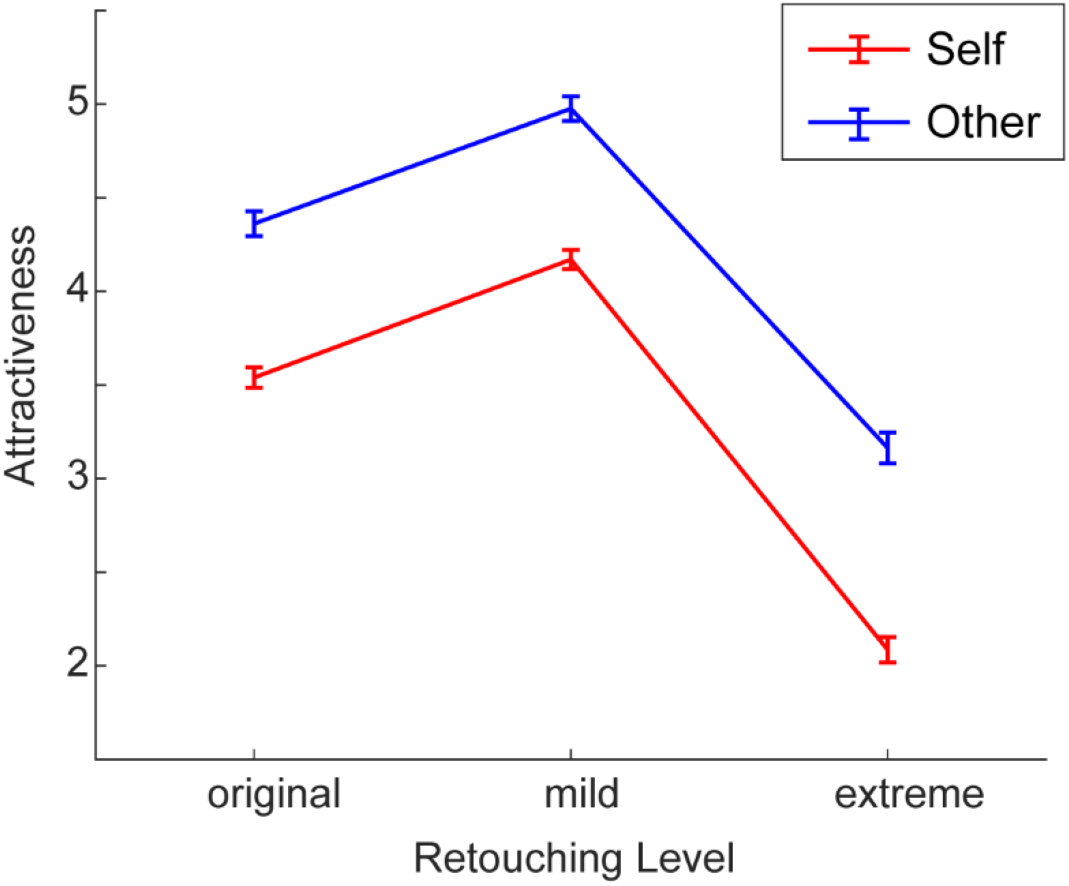
Comparison of attractiveness scores between the 3 retouching levels. Red and blue lines represent the score for self-face and other-face, respectively. The error bar represents a standard error.

### Brain activity correlated with attractiveness score

To identify brain regions that show changes in activity correlated with facial attractiveness, we conducted a parametric modulation analysis using the participants’ attractiveness rating for each face stimuli for whole brain. The neural activity in the NA, mPFC, Anterior insula (AI), and precuneus showed a significant positive correlation with attractiveness scores (Table 1, Fig. 3A). On the other hand, the ROI-based analysis revealed that the neural activity in the amygdala showed a significant negative correlation with them (Table 1, Fig. 3B), although it did not survive in the whole brain exploratory analysis. Since the participants responded using a score of 1-4 with the left hand and a score of 5-8 with the right hand, the right cerebellum and left motor cortex showed activation in positive association with attractiveness scores, and the left cerebellum and right motor cortex did activation in negative association with them (Table 1). We analyzed brain regions showing changes in activity positively correlated with facial attractiveness for self-face and other-face separately. The results showed that bilateral NA consistently increased activity in association with facial attractiveness for both faces (see supplementary figure 2).

**Fig. 3.**
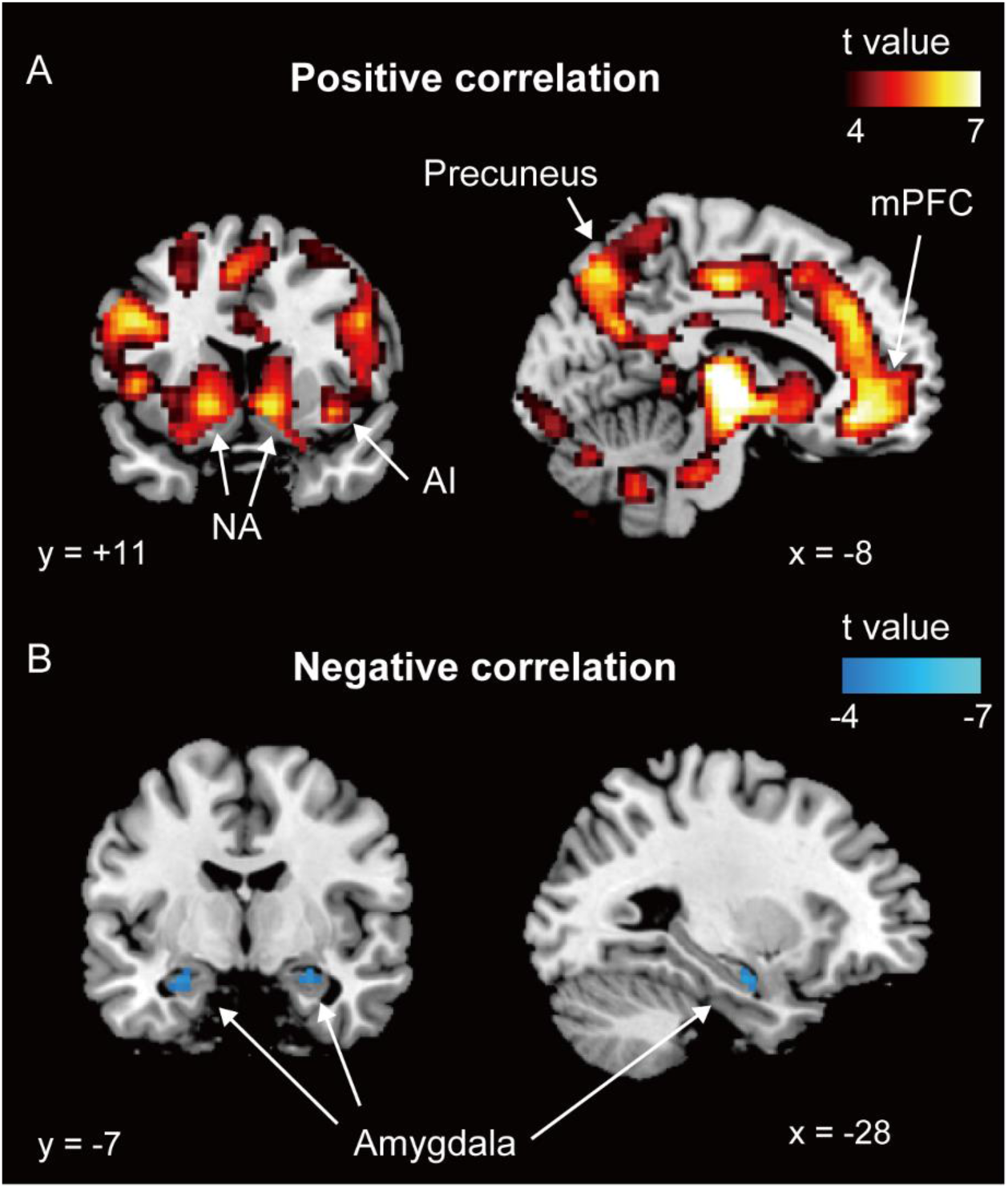
Brain regions exhibiting a significant correlation with facial attractiveness by the parametric modulation analysis. (A) Brain regions showing positive correlation with facial attractiveness (FWE-corrected threshold p < 0.05 for cluster level with a voxel-level uncorrected height threshold p < 0.001). (B) Brain regions showing negative correlation with facial attractiveness (FEW-corrected threshold p<0.05 with small volume correction for cluster level with a voxel-level uncorrected height threshold p < 0.001). The color bars represent voxel-level t-values.

**Table 1.**
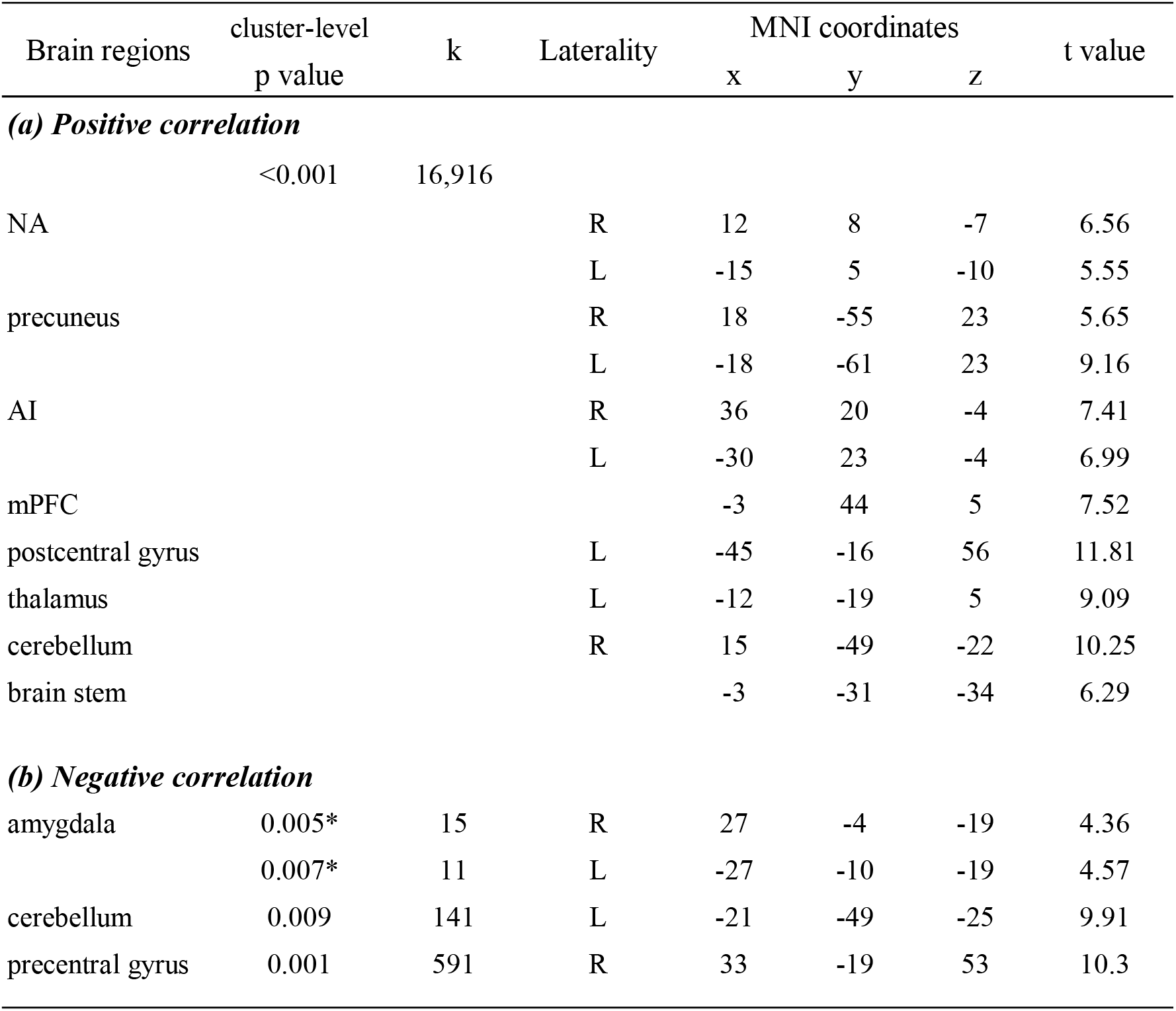
Brain regions showing activity changes associated with facial attractiveness by the parametric modulation analysis. The cluster-level statistics uses the FWE-corrected threshold p < 0.05, while * was applied the FWE corrected cluster-level threshold of p < 0.05 with small volume correction. The t values represent voxel-level uncorrected statistics (p < 0.001).

| Brain regions | cluster-level | k | Laterality | MNI coordinates |  |  | t value |
| --- | --- | --- | --- | --- | --- | --- | --- |
|  | p value |  |  | x | y | z |  |
| *(a) Positive correlation* |  |  |  |  |  |  |  |
|  | <0.001 | 16,916 |  |  |  |  |  |
| NA |  |  | R | 12 | 8 | -7 | 6.56 |
|  |  |  | L | -15 | 5 | -10 | 5.55 |
| precuneus |  |  | R | 18 | -55 | 23 | 5.65 |
|  |  |  | L | -18 | -61 | 23 | 9.16 |
| AI |  |  | R | 36 | 20 | -4 | 7.41 |
|  |  |  | L | -30 | 23 | -4 | 6.99 |
| mPFC |  |  |  | -3 | 44 | 5 | 7.52 |
| postcentral gyrus |  |  | L | -45 | -16 | 56 | 11.81 |
| thalamus |  |  | L | -12 | -19 | 5 | 9.09 |
| cerebellum |  |  | R | 15 | -49 | -22 | 10.25 |
| brain stem |  |  |  | -3 | -31 | -34 | 6.29 |
| *(b) Negative correlation* |  |  |  |  |  |  |  |
| amygdala | 0.005* | 15 | R | 27 | -4 | -19 | 4.36 |
|  | 0.007* | 11 | L | -27 | -10 | -19 | 4.57 |
| cerebellum | 0.009 | 141 | L | -21 | -49 | -25 | 9.91 |
| precentral gyrus | 0.001 | 591 | R | 33 | -19 | 53 | 10.3 |

We further compared the time courses of the BOLD signal of the five brain regions (NA, amygdala, mPFC, AI, precuneus) between the retouching levels. The NA showed an increase of the BOLD signal with a peak at 5 s after the stimulus onset of the self-face with mild retouching, whereas it showed a decrease of the BOLD signal in response to the self-face with extreme retouching and the other-face (Fig. 4A). Correspondingly, the mean beta value was positive for the self-face with both mild and no retouching but negative for the self-face with extreme retouching and the other-face (Fig.4B). Two-way ANOVA with factors of face type and retouching level confirmed a significant interaction (F_2,64_ = 6.6, *p* = 0.0024). In the post hoc analysis, the beta values of the NA for the self-face with mild retouching and no retouching was significantly higher than that of the self-face with extreme retouching (mild retouching vs. extreme retouching t_128_ = 5.6, p < 0.00001; no retouching vs. extreme retouching t_128_ = 4.2, p < 0.0001). No significant difference was detected between no retouching versus mild retouching in the self-face condition (t_128_ =1.4, p < 0.2). In addition, the beta values for the cases of self-face with no retouching and mild retouching were significantly higher than those for the cases of other-face with no retouching and mild retouching, respectively (no retouching F_96_ = 13.8, p = 0.0003, mild retouching F_96_ = 16.8, p = 0.0001).

**Fig. 4.**
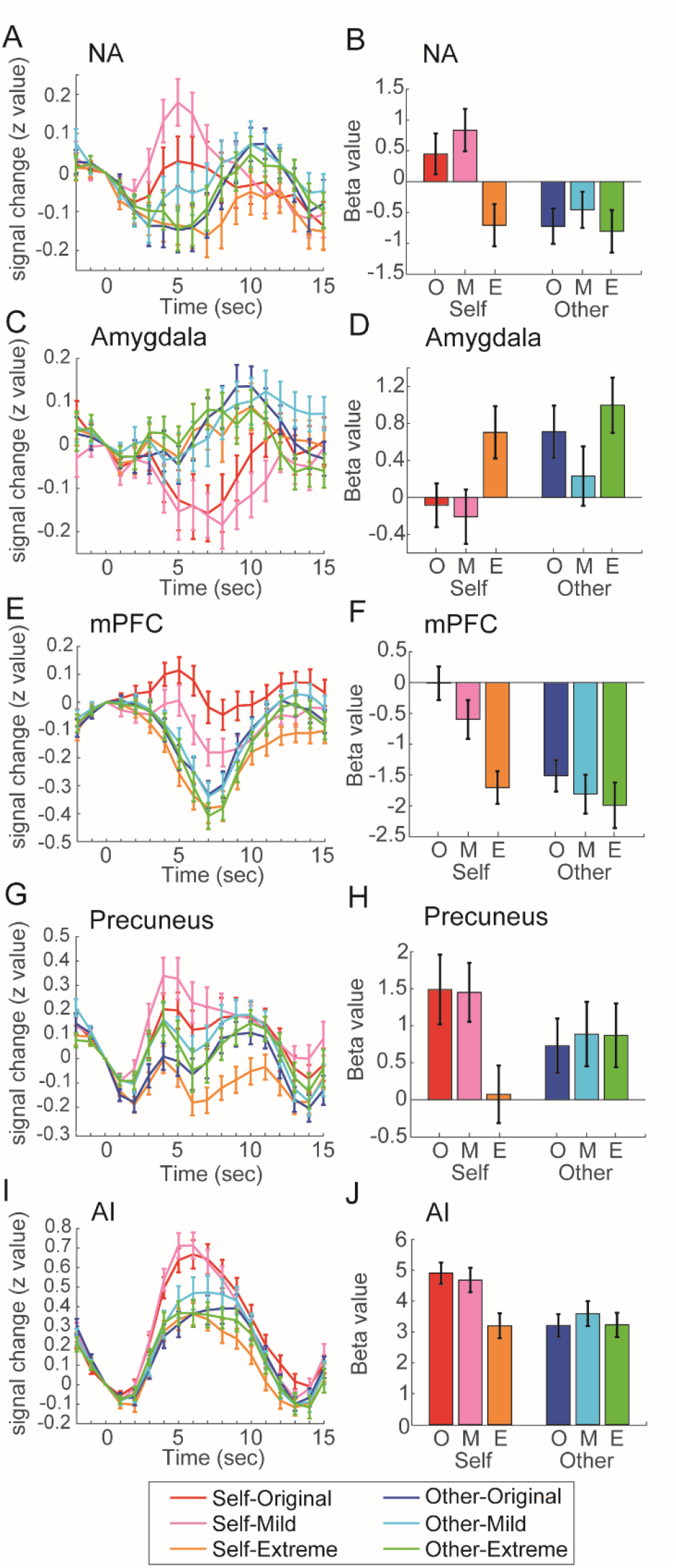
ROI analysis of BOLD signal changes and beta value. Mean time courses of signal intensity (A,C,E,G,I) and mean of beta value (B,D,F,H,J) in each ROI in response to retouching condition for self and other faces. In the horizontal axis of the right side panels, “O”, “M”, “E” represents original, mild and extreme retouching, respectively. The error bars represent standard error. NA: nucleus accumbens; mPFC: medial prefrontal cortex; AI: anterior insula.

The amygdala showed an opposite response pattern to the NA (Fig 4C,D). The BOLD signal gradually increased with a peak at 10 s in response to the other-face and the self-face with an extreme retouching. In contrast, it rapidly decreased with a peak at 7 s in response to the self-face with either no retouching or mild retouching. The ANOVA detected significant main effects of retouching level (F_2,64_ = 12.5, p < 0.0001) and face type (F_1,32_ = 18.6, p < 0.0001) but not significant interaction of the beta value (F_2,64_=1.3, p = 0.3). The post-hoc test confirmed that the amygdala showed significantly higher activation in response to the extreme retouching than to both mild and no retouching (extreme vs. none t_64_=3.2, p < 0.002; extreme vs. mild t_64_=4.9, p < 0.00001).

The mPFC and precuneus, which belong to the default mode network that decreases the brain activity to the external visual stimuli, both consistently showed a tendency for the BOLD signal to decrease in response to the facial stimuli except in the cases self-face with mild or no filter (Fig.4E&G). However, the temporal dynamics of the BOLD signal were different between the mPFC and precuneus. The BOLD signal in the mPFC reached a minimum at 7 s after the face onset (Fig.4E), whereas in the precuneus the signal reached a minimum twice, 4 s and 10 s after the face onset (Fig.4G). The two-way ANOVA confirmed a significant interaction of the beta value in both brain regions (mPFC F_2,64_ = 5.3, p<0.007, precuneus F_2,64_=8.1, p < 0.0007). The beta values of the self-face with mild or no retouching was significantly higher than that of the self-face with extreme retouching (mPFC, none vs. extreme t_128_ = 5.9, p<0.00001; mild vs. extreme t_128_ = 3.8, p<0.0002; precuneus, none vs. extreme t_128_=4.9, p < 0.00001; mild vs. extreme t_128_=4.8, p < 0.00001), whereas the other-face condition had no significant difference among retouching s (Fig.4F,H).

In contrast to the default mode network, the AI showed hemodynamic responses which peaked at 5s after stimulus onset for all conditions (Fig. 4I). Two-way ANOVA detected significant interaction of the beta value in the AI (F_2,64_=12.7, p < 0.00001). The post hoc analysis revealed that the AI showed a greater increase in brain activity in response to cases of self-face with no filter or mild retouching than cases of self-face with extreme retouching (none vs. extreme t_128_=6.9, p < 0.00001; mild vs. extreme t_128_=6.0, p < 0.00001), whereas no significant difference in brain activity was observed between retouching levels in the other-face condition (Fig 4J).

We further investigated the flow of information between these five brain regions by the MVGC method, which analyze the causal relationships between multiple time series data. As shown in Fig5, information is transmitted from the amygdala to the NA and the precuneus. The NA also receives information from the precuneus, AI and mPFC.

**Fig. 5.**
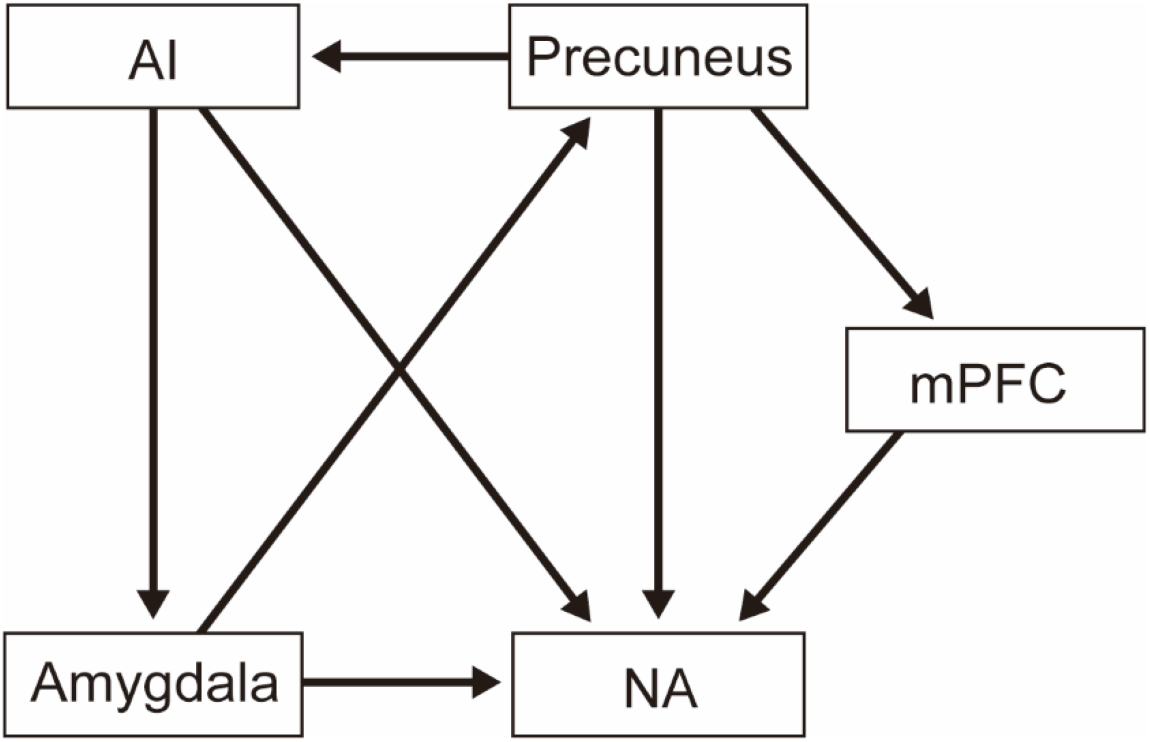
Functional connectivity between brain regions related to facial attractiveness. The arrows indicate the direction of information transfer revealed by the MVGC analysis (p < 0.05, multiple comparisons were corrected by Bonferroni method). NA: nucleus accumbens; mPFC: medial prefrontal cortex; AI: anterior insula.

### Self vs other-face comparison

We further investigated differences in brain activity between self-face and other-face conditions for each retouching level. As shown in Fig.6A and Table 2, cases of the original self-face induced significantly greater activation than cases of the original other-face in broad brain regions including the precentral gyrus, middle temporal gyrus (MTG), inferior frontal gyrus (IFG), supramarginal gyrus (SMG), ACC, PCC, and subcortical regions. The same brain regions also exhibited greater activation to in cases of the self-face with mild filter than cases of other-face with mild filter, but the activation areas were smaller than the original face condition (Fig.6B, Table 2). In the extreme retouching condition, no significant difference in activation between self and other faces was found in any brain area. On the other hand, the ROI based statistical analysis revealed that the amygdala exhibited greater activation to the other-face than the self-face across all filter levels (Fig.6C, Table 2), although it did not survive in the whole brain exploratory analysis.

**Fig. 6.**
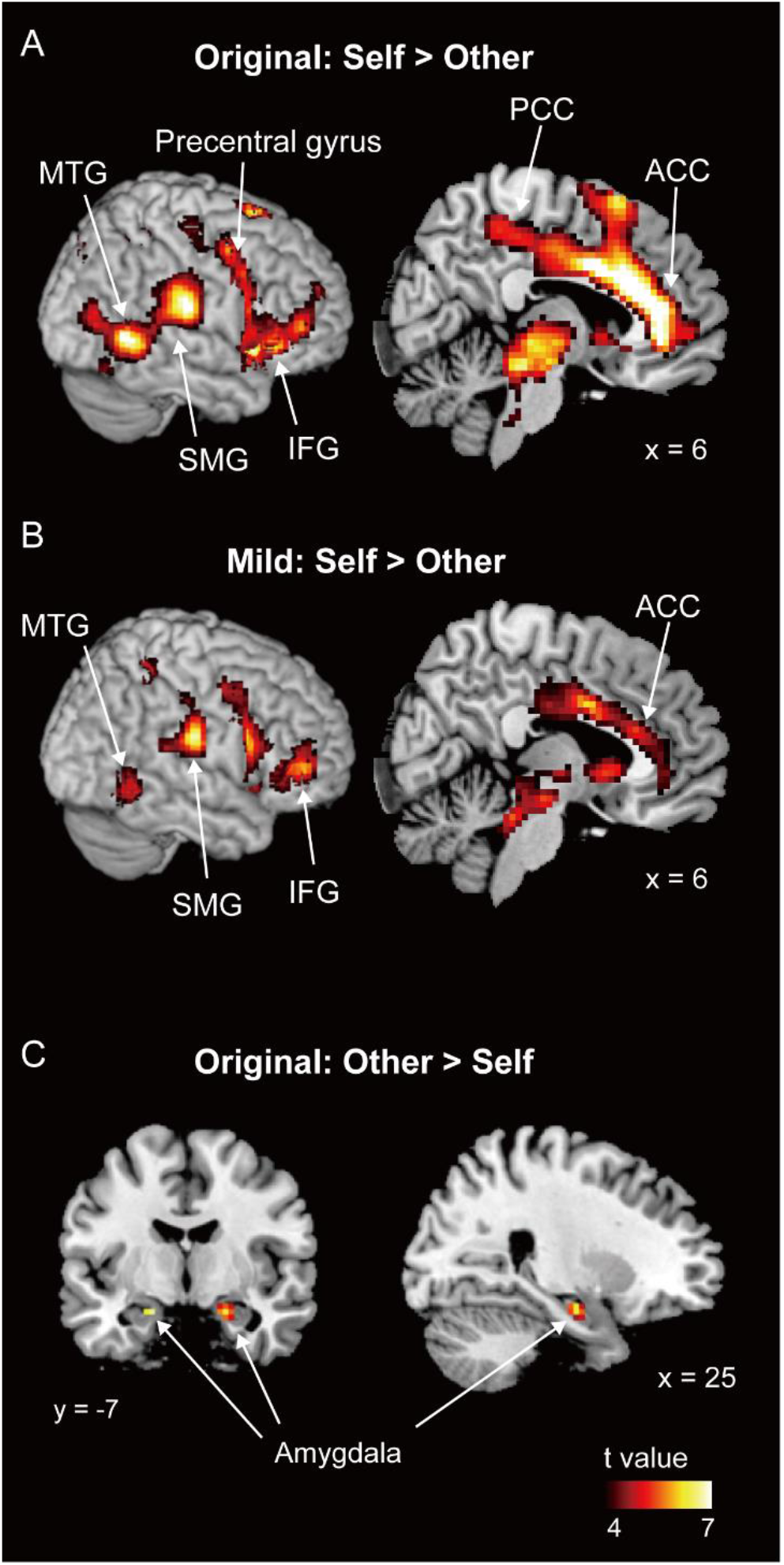
Comparison of brain activation between self-face versus other-face conditions. Brain regions showing greater activation for self-face condition than other-face condition without any retouching (A), with a mild retouching (B) (FWE-corrected threshold p < 0.05 for cluster level with a voxel-level uncorrected height threshold p < 0.001). (C) Brain regions showing greater activation for other-face condition than self-face condition without any retouching (FEW-corrected threshold p < 0.05 with small volume correction for cluster level with a voxel-level uncorrected height threshold p < 0.001). The color bars represent voxel-level t-values. SMG: supramarginal gyrus; MTG: middle temporal gyrus; IFG: inferior frontal gyrus; PCC: posterior cingulate cortex; ACC: anterior cingulate cortex;

**Table 2.** Brain regions showing a significant difference between the self-face and the other-face for each filter level. The cluster-level statistics uses the FWE-corrected threshold p < 0.05. * was applied the FWE corrected cluster-level threshold of p < 0.05 with the small volume correction. The t values represent voxel-level uncorrected statistics (p < 0.001).

| Brain regions | cluster-level | k | Laterality | MNI coordinates |  |  | t value |
| --- | --- | --- | --- | --- | --- | --- | --- |
|  | p value |  |  | x | y | z |  |
| *self > other* |  |  |  |  |  |  |  |
| *(a) original (no filter)* |  |  |  |  |  |  |  |
|  | <0.001 | 9,732 |  |  |  |  |  |
| IFG |  |  | R | 48 | 29 | 5 | 6.55 |
| precentral gyrus |  |  | R | 45 | 2 | 50 | 5.56 |
| MTG |  |  | R | 60 | -55 | 5 | 7.46 |
|  |  |  | L | -48 | -58 | 17 | 7.33 |
| SMG |  |  | R | 63 | -22 | 23 | 7.29 |
|  |  |  | L | -63 | -37 | 32 | 6.23 |
| ACC |  |  | R/L | 0 | 26 | 23 | 9.57 |
| PCC |  |  | R/L | 12 | -31 | 41 | 6.81 |
| anterior insula |  |  | R | 36 | 11 | -7 | 8.05 |
|  |  |  | L | -39 | 11 | -7 | 6.44 |
| brain stem |  |  | R/L | 9 | -25 | -13 | 7.08 |
| *(b) mild filter* |  |  |  |  |  |  |  |
|  | <0.001 | 1,357 |  |  |  |  |  |
| IFG |  |  | R | 48 | 44 | 5 | 5.7 |
| anterior insula |  |  | R | 39 | 8 | -7 | 7.39 |
|  | <0.001 | 663 |  |  |  |  |  |
| MTG |  |  | L | -57 | -52 | -1 | 5.19 |
| SMG |  |  | L | -45 | -52 | 23 | 4.74 |
|  | <0.001 | 585 |  |  |  |  |  |
| SMG |  |  | R | 63 | -19 | 29 | 6.73 |
| SPL |  |  | R | 27 | -70 | 38 | 4.72 |
| brain stem | 0.001 | 874 | R/L | -6 | -34 | -22 | 5.36 |
| MTG | 0.001 | 387 | R | 51 | -49 | -1 | 5.53 |
| ACC | 0.001 | 910 | R/L | 3 | 5 | 32 | 6.15 |
| anterior insula | 0.001 | 524 | L | -39 | 5 | -7 | 5.24 |
| *other > self* |  |  |  |  |  |  |  |
| *(c) original (no filter)* |  |  |  |  |  |  |  |
| amygdala | 0.005* | 17 | R | 21 | -7 | -19 | 6.29 |
|  | 0.005* | 7 | L | -24 | -7 | -19 | 5.11 |
| precentral gyrus | 0.021 | 106 | L | -36 | -19 | 53 | 4.41 |
| *(d) mild filter* |  |  |  |  |  |  |  |
| amygdala | 0.01* | 6 | R | 21 | -7 | -16 | 4.52 |
|  | 0.01* | 7 | L | -21 | -7 | -16 | 6 |
| *(e) extreme filter* |  |  |  |  |  |  |  |
| amygdala | 0.014* | 3 | R | 21 | -7 | -19 | 3.9 |
| posterior insula | 0.001 | 447 | L | -45 | -22 | 11 | 6.45 |
| precentral gyrus | 0.001 | 1,465 | L | -15 | -16 | 62 | 6.21 |
| lingual gyrus | 0.001 | 921 | R/L | 6 | -85 | -4 | 5.99 |
| fusiform gyrus | 0.015 | 131 | R | 33 | -37 | -22 | 5.17 |
| MTG | 0.004 | 180 | R | 63 | -7 | -19 | 4.77 |

## Discussion

Our perception of facial attractiveness changes in an inverted-U shaped manner depending on the level of beauty retouching applied, such that mildly-retouched faces look more attractive than the original face but extremely-retouched faces look less attractive than the original (Nakano & Uesugi, 2020). The present study replicated this phenomenon and examined its neural correlates via fMRI. We revealed that the activity of the reward circuits that include the NA and mPFC showed positive correlation with facial attractiveness. These regions showed activation in response to positive modulation of attractiveness in the self-face condition but not in the other-face condition. In contrast, activity in the amygdala, which is the center of negative value coding (Adolphs et al., 1999; LeDoux, 2000), showed a negative correlation with facial attractiveness. In both face types, the extreme beauty retouching induced greater activation in the amygdala than was seen in either the mild-retouching or no-retouching condition. The causal-relationship analysis using the MVGC method revealed that amygdala has a causal effect on the activity of the NA. Consistently, a previous anatomical study reported that the amygdala has strong projections to the NA (McDonald, 1991). Considering these results, the NA receives inhibitory modulation by the amygdala. These results suggest that the dynamic interactions between the NA and amygdala underlie the dramatic changes seen in our perception of facial attractiveness based on levels of beauty retouching.

One of the main findings in the present study is that the NA, the core region of our neural reward system (Covey & Cheer, 2019), showed increased activation only in cases of a self-face with mild beauty retouching. At the behavioral level, the positive modulation of facial attractiveness by beauty retouching was consistently observed in both self and other face conditions. Additionally, despite participants consistently rating others’ faces as being more attractive than their own, the positive valence system reacted only in the self-face and not the other-face condition. This raises the possibility that the neural mechanisms for subjective value coding of beauty are isolated from those for objective beauty evaluation. Correspondingly, a previous study reported that, although males rated pictures of both beautiful males and females as attractive, their NA exhibited activation only towards beautiful female faces (Aharon et al., 2001). In addition, the positive modulation of attractiveness by contrast effect induced activation in the reward-related brain regions for the self-face but not for the other-face condition (Oikawa et al., 2012). These results indicate that facial beauty can be evaluated even if the reward system does not work, and that the reward system works only for specific facial information. Since reward-value coding is important for action selection (Dayan & Balleine, 2002), the reward system reacts only to facial information related to selecting a romantic partner or to improving oneself.

It is also worth noting that the brain regions corresponding to positive modulation of facial attractiveness overlap with the regions involved in self-related processing. This layout might be closely related to our observation that the reward system was activated only in the self-face condition. It is a well-known fact that several brain regions exhibit self-specific activation. Specifically, regarding facial information, the right lateralized cortical regions show self-face specific activation (Sugiura, 2015; Uddin, Kaplan, Molnar-Szakacs, Zaidel, & Iacoboni, 2005). This is replicated in the present study, in which the right IFG, bilateral SMG, and MTG showed greater activation in the self-face than the other-face condition. In addition to self-face specific activation, the cortical midline areas exhibit specific activation towards various types of self-related information (Northoff et al., 2006). Consistently, the present study reported that these cortical midline areas (ACC and PCC) showed greater activation towards cases of self-face than other-face. Furthermore, the medial frontal area was also involved in coding positive modulation of facial attractiveness. These results suggest that the cortical networks involved in recognizing one’s own face interact with the networks involved in self-referential processing and value-coding. It might induce self-face specific activation in the reward system. Interestingly, the images of self-face with an extreme retouching did not activate the cortical midline areas, the right IFG and the bilateral temporal areas. This suggests that the brain no longer recognizes a face that has been altered significantly as its own face.

Another important finding of the present study is that the amygdala showed greater activation towards the extremely retouched faces than towards the mildly retouched or no retouched faces. Faces with unnatural local-global balance due to extreme retouching (e.g. extremely large eyes on very thin chin) look human-like but do not look like real humans. Such subtle differences between real humans and human-like artifacts induce strong eerie feelings in people: this is known as the “uncanny valley phenomenon” (Mori et al., 2012). A previous study examined the neural correlates of the Thatcher effect that the upright face with inverted eyes and mouth both looks very creepy, and reported strong activation in the bilateral amygdala to the creepy face (Rotshtein et al., 2001). Another previous study also reported that large eyes induced the amygdala’s response(Whalen et al., 2004). Considering these facts, the amygdala response to the extremely retouched face may not only be related to low attractiveness, but also is involved in the detection of large-eyed, abnormal faces and the generation of an eerie sensation. As described above, since positive modulation by beauty retouching activates the reward system, facial retouching behaviors using beauty filter app become reinforced. In fact, young women prefer to apply stronger beauty filters on their own face than on others’ faces (Nakano & Uesugi, 2020). Conversely, the amygdala induces negative reactions towards excessively retouched faces, which helps suppress the reinforced behavior of retouching. If this neural system of the uncanny valley phenomenon did not exist, people might have started looking increasingly artificial to the point of practically zombie-like from excessive use of cosmetics.

The amygdala also showed activation in the other-face condition regardless of retouching level in the present study. Since all pictures in the other-face condition were of persons unknown to the participants, we can infer that poor affinity towards unknown faces might also activate the amygdala, regardless of attractiveness (Schwartz et al., 2003). It is also worth noting that while the amygdala gradually showed increased activity in response to unknown faces, it displayed a rapid decrease in activity in response to self-face with either mild-retouching or no-retouching (see Fig.4C). Recent studies in rodents and non-human primates also revealed populations of valence-selective neurons in their amygdala such that some neurons, excited by a negative cue, show inhibition in the presence of the reward cue, and vice versa (Janak & Tye, 2015). These results suggest that the same region of the amygdala is involved in coding both negative and positive values by controlling neural activity through increase or inhibition.

### Conclusions

The present study revealed the dynamic interactions between the NA and amygdala underlies the non-linear change in perceived facial attractiveness depending on the level of beauty retouching. This NA reacted specific to the positive modulation of self-face, suggesting that self-related positive value is critical for action selection compared to other-related positive value. On the other hand, the amygdala plays a central role in generation of the “uncanny valley” phenomenon, which might prevent us from excessive dependence of makeup, facial retouching or cosmetic surgery.

## Acknowledgements

We thank for Malini V. Nair for editing for English language of the manuscript. This work was supported by a Grant-in-Aid 18H04084 awarded to TN from the Ministry of Education, Culture, Sports, Science and Technology, Japan, as well as a PRESTO grant, “The Future of Humans and Interactions” #30227, awarded to TN from the Japan Science and Technology Agency, Japan.

## References

Adolphs, R., Russell, J. A., & Tranel, D. (1999). A role for the human amygdala in recognizing emotional arousal from unplesant stimuli. Psychological Sciene, 10(2), 167–171.

Aharon, I., Etcoff, N., Ariely, D., Chabris, C. F., O’Connor, E., & Breiter, H. C. (2001). Beautiful faces have variable reward value: fMRI and behavioral evidence. Neuron, 32(3), 537–551. doi:10.1016/s0896-6273(01)00491-3

Covey, D. P., & Cheer, J. F. (2019). Accumbal Dopamine Release Tracks the Expectation of Dopamine Neuron-Mediated Reinforcement. Cell Rep, 27(2), 481–490 e483. doi:10.1016/j.celrep.2019.03.055

Dayan, P., & Balleine, B. W. (2002). Reward, motivation, and reinforcement learning. Neuron, 36(2), 285–298. doi:10.1016/s0896-6273(02)00963-7

Feng, S., Wang, X., Wang, Q., Fang, J., Wu, Y., Yi, L., & Wei, K. (2018). The uncanny valley effect in typically developing children and its absence in children with autism spectrum disorders. PLoS One, 13(11), e0206343. doi:10.1371/journal.pone.0206343

Forbes, T. R. (1977). Why is it called ‘beautiful lady’? A note on belladonna. Bull NY Acad Med, 53(4), 403–406.

Janak, P. H., & Tye, K. M. (2015). From circuits to behaviour in the amygdala. Nature, 517(7534), 284–292. doi:10.1038/nature14188

Kee, E., & Farid, H. (2011). A perceptual metric for photo retouching. Proc Natl Acad Sci U S A, 108(50), 19907–19912. doi:10.1073/pnas.1110747108

LeDoux, J. E. (2000). Emotion circuits in the brain. Annu Rev Neurosci, 23, 155–184. doi:10.1146/annurev.neuro.23.1.155

McDonald, A. J. (1991). TOPOGRAPHICAL ORGANIZATION OF AMYGDALOID PROJECTIONS TO THE CAUDATOPUTAMEN, NUCLEUS ACCUMBENS, AND RELATED STRIATAL-LIKE AREAS OF THE RAT BRAIN. Neuroscience, 44, 15–33.

Mori, M., MacDorman, K., & Kageki, N. (2012). The Uncanny Valley [From the Field]. IEEE Robotics & Automation Magazine, 19(2), 98–100. doi:10.1109/mra.2012.2192811

Motoki, K., Sugiura, M., Takeuchi, H., Kotozaki, Y., Nakagawa, S., Yokoyama, R., & Kawashima, R. (2016). Are Plasma Oxytocin and Vasopressin Levels Reflective of Amygdala Activation during the Processing of Negative Emotions? A Preliminary Study. Frontiers in Psychology, 7(480). doi:10.3389/fpsyg.2016.00480

Nakano, T., & Uesugi, Y. (2020). Risk Factors Leading to Preference for Extreme Facial Retouching. Cyberpsychol Behav Soc Netw, 23(1), 52–59. doi:10.1089/cyber.2019.0545

Northoff, G., Heinzel, A., de Greck, M., Bermpohl, F., Dobrowolny, H., & Panksepp, J. (2006). Self-referential processing in our brain--a meta-analysis of imaging studies on the self. Neuroimage, 31(1), 440–457. doi:10.1016/j.neuroimage.2005.12.002

O’Doherty, J., Winston, J., Critchley, H., Perrett, D., Burt, D. M., & Dolan, R. J. (2003). Beauty in a smile: the role of medial orbitofrontal cortex in facial attractiveness. Neuropsychologia, 41(2), 147–155. doi:10.1016/s0028-3932(02)00145-8

Oikawa, H., Sugiura, M., Sekiguchi, A., Tsukiura, T., Miyauchi, C. M., Hashimoto, T., . . . Kawashima, R. (2012). Self-face evaluation and self-esteem in young females: an fMRI study using contrast effect. Neuroimage, 59(4), 3668–3676. doi:10.1016/j.neuroimage.2011.10.098

Rotshtein, P., Malach, R., Hadar, U., Graif, M., & Hendler, T. (2001). Feeling or Features: Different Sensitivity to Emotion in High-Order Visual Cortex and Amygdala. Neuron, 32, 747–757.

Schienle, A., Schafer, A., Stark, R., Walter, B., & Vaitl, D. (2005). Relationship between disgust sensitivity, trait anxiety and brain activity during disgust induction. Neuropsychobiology, 51(2), 86–92. doi:10.1159/000084165

Schwartz, C. E., Wright, C. I., Shin, L. M., Kagan, J., Whalen, P. J., McMullin, K. G., & Rauch, S. L. (2003). Differential amygdalar response to novel versus newly familiar neutral faces: a functional MRI probe developed for studying inhibited temperament. Biol Psychiatry, 53(10), 854–862. doi:10.1016/s0006-3223(02)01906-6

Sugiura, M. (2015). Three faces of self-face recognition: potential for a multi-dimensional diagnostic tool. Neurosci Res, 90, 56–64. doi:10.1016/j.neures.2014.10.002

Thompson, P. (1980). Margaret Thatcher: a new illusion. Perception, 9(4), 483–484. doi:10.1068/p090483

Uddin, L. Q., Kaplan, J. T., Molnar-Szakacs, I., Zaidel, E., & Iacoboni, M. (2005). Self-face recognition activates a frontoparietal “mirror” network in the right hemisphere: an event-related fMRI study. Neuroimage, 25(3), 926–935. doi:10.1016/j.neuroimage.2004.12.018

Ueno, A., Ito, A., Kawasaki, I., Kawachi, Y., Yoshida, K., Murakami, Y., . . . Fujii, T. (2014). Neural activity associated with enhanced facial attractiveness by cosmetics use. Neurosci Lett, 566, 142–146. doi:10.1016/j.neulet.2014.02.047

Vuilleumier, P., Richardson, M. P., Armony, J. L., Driver, J., & Dolan, R. J. (2004). Distant influences of amygdala lesion on visual cortical activation during emotional face processing. Nat Neurosci, 7(11), 1271–1278. doi:10.1038/nn1341

Whalen, P. J., Kagan, J., Cook, R. G., Davis, F. C., Kim, H., Polis, S., . . . Johnstone, T. (2004). Human amygdala responsivity to masked fearful eye whites. Science, 306(5704), 2061. doi:10.1126/science.1103617

Winston, J. S., O’Doherty, J., Kilner, J. M., Perrett, D. I., & Dolan, R. J. (2007). Brain systems for assessing facial attractiveness. Neuropsychologia, 45(1), 195–206. doi:10.1016/j.neuropsychologia.2006.05.009

