## Supplemental Figures for "Neural correlates of beauty retouching to enhance attractiveness of self-depictions in women"

Supplementary Figure 1.

Section images of the implicit mask for group-analysis. Brain regions covered by the implicit mask were plotted in white and superimposed on structural images.

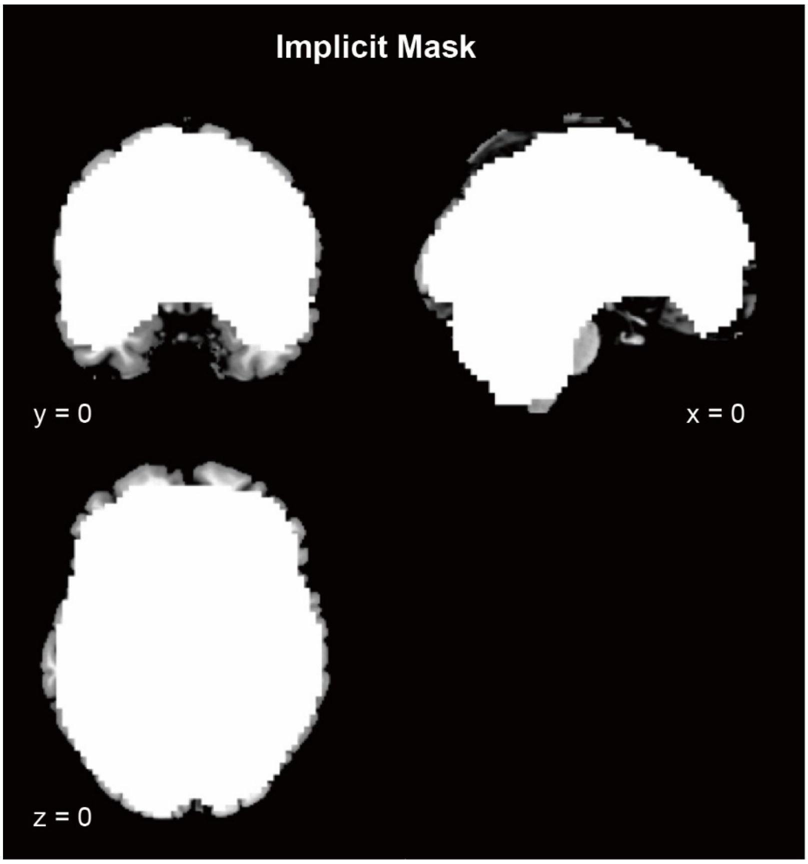

The Supplementary Figure. 2.

Brain regions showing positive correlation with facial attractiveness in self-face (A) and other-face (B) conditions by parametric modulation analysis. NA: nucleus accumbens.

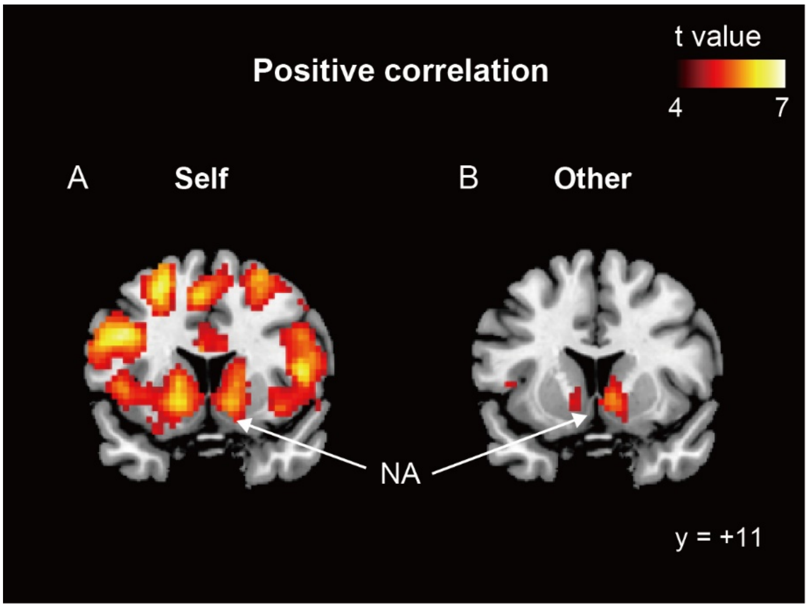
